# MSIsensor-pro: fast, accurate and matched-normal-sample-free detection of microsatellite instability

**DOI:** 10.1101/2020.01.08.899633

**Authors:** Peng Jia, Xiaofei Yang, Li Guo, Bowen Liu, Jiadong Lin, Hao Liang, Jianyong Sun, Chengsheng Zhang, Kai Ye

## Abstract

We developed MSIsensor-pro (https://github.com/xjtu-omics/msisensor-pro), an open-source single sample microsatellite instability (MSI) scoring method for research and clinical applications. MSIsensor-pro introduces a multinomial distribution model to quantify polymerase slippages for each tumor sample and a discriminative sites selection method to enable MSI detection without matched normal samples. For samples of various sequencing depths and tumor purities, MSIsensor-pro significantly outperformed the current leading methods in terms of both accuracy and computational cost.

## MAIN TEXT

Microsatellite instability (MSI) is a form of hypermutation of the microsatellites in malignancies due to a deficient DNA mismatch repair (MMR) system^1–3^. Significant proportions of tumor samples with MSI status are observed in colorectal cancer (CRC), stomach adenocarcinoma (STAD) and uterine corpus endometrial carcinoma (UCEC)^4,5^. Given that MSI is an important molecular phenotype of cancers and a key biomarker for cancer immunotherapy^6–8^, two gold standards, MSI-PCR and MSI-IHC, are widely used for clinical MSI identification^9^. However, both methods are laborious, time-consuming and expensive^9,10^. Recently, several next-generation-sequencing (NGS)-based methods have been developed, which show much better time and cost efficiency and are highly consistent with the gold standards^3–5,11–15^. For instance, MSIsensor^11^, an FDA-approved MSI detection solution of MSK-IMPACT^16^, achieved 99.4% concordance and high sensitivity^17^. However, these NGS methods do have limitations, such as requiring matched normal samples as control (sometimes inaccessible), being computational expensive, and being affected by low sequencing depths and low tumor purities^9^.

A hallmark of MSI is the enrichment of insertions or deletions in microsatellite regions initiated by polymerase slippages^2,18^ (Supplementary Fig. 1), an enzymatic process that we argue is described using a multinomial distribution (MND) model (Supplementary Fig. 2), providing promising improvements of MSI detection efficacy on NGS data. Here, we report a novel MSI calling method, MSIsensor-pro, which addresses the limitations of current NGS-based MSI detection tools by applying MND model to capture the intrinsic properties of polymerase slippages in a single sample. We demonstrate that MSIsensor-pro is an ultrafast, accurate and normal sample-free MSI calling method. Moreover, it outperforms all current MSI detection methods and is robust for samples with various sequencing depths, tumor purities and target sequencing regions.

To quantitatively describe the polymerase slippages present in a single sample, we first examined the allele length distributions of 27,200 microsatellites in 1,532 normal samples from the Cancer Genome Atlas (TCGA)^19^ (Supplementary Table 1,2, Methods). The distributions flattened (the variances became larger and the modes deviated from expectation) with the increase in the repeat length of microsatellites on the reference genome (Fig. 1a), suggesting that the polymerase slippage was an iterative process. We argue that polymerase slippages are independently accumulative in the DNA replication process and could be modeled by a MND model. Here, we used *p* and *q* to denote the probabilities of hysteresis synthesis (causing deletions) and presynthesis (causing insertions), respectively, for each replication unit (Supplementary Fig. 2). We next estimated the parameters *p* and *q* of each microsatellite to quantify the polymerase slippage in a given allele length distribution.

To explore the characteristics of parameter *p* and *q* in MND model, we applied MND model to 1,532 TCGA normal samples. We totally obtained 11,666 microsatellites with sufficient read coverage (>20) in more than half of the samples for subsequent study (Supplementary Table 1-2, Methods). We found that the average probability of hysteresis synthesis, *p*, is significantly larger (*P*-value <0.05, Wilcoxon rank-sum test) than the average probability of presynthesis, *q* (Supplementary Fig. 3) in these sites, indicating that polymerase slippages tend to cause more deletions than insertions at microsatellites, confirming previous reports^4,18^. To evaluate the power of our MND model on describing the polymerase slippages in DNA replication, we simulated the allele length distributions at each microsatellite site with their corresponding computed *p* and *q* values and compared them with the observed ones from sequencing data. We found that the simulated allele length distributions were consistent with the observed ones at 91.97% microsatellites and the similarities of two distributions decreased with increasing repeat length (Fig. 1b, Supplementary Figs. 4, 5 and Methods), confirming the MND model is capable of describing the polymerase slippages at microsatellite sites..

Based on the MND model, we developed a method called MSIsensor-pro to detect MSI. We applied our MND model to 1,532 TCGA tumor samples with clinical MSI status and obtained their *p* and *q* values for each microsatellite. We found that the MSI samples have significantly larger *p* values than do MSS samples (*P*-value < 2×10^16^), while *q* values in the MSI and MSS samples are not discriminative (Fig. 1c, d and Supplementary Figs. 6-9). Thus, it is conceivable that the higher incidence of polymerase slippages, and therefore, the greater instability of microsatellites in MSI instead of MSS, is attributed to more deletions, rather than insertions. Therefore, the parameter *p* evaluates the stability of each microsatellite site. MSIsensor-pro classifies the *i*-th microsatellite as unstable when its *p* is larger than *μ_i_+3σ_i_*, in which *μ_i_* and *σ_i_* are the mean and standard deviation, respectively, of *p* in 1,532 normal samples at the *i-th* microsatellite. The fraction of unstable sites in a given microsatellite set is used to score MSI in a tumor sample (Supplementary Fig. 10 and Methods).

To assess the performance of MSIsensor-pro in terms of the accuracy and computational cost, we compared this method against MSIsensor^11^, mantis^13^ and mSINGS^12^; the first two methods require tumor-normal-paired samples and the last method requires tumor-only samples (Supplementary Tables 1-2 and Supplementary Note). First, we applied MSIsensor-pro to 1,532 TCGA tumor samples based on 11,666 preselected microsatellites to detect MSI and then compared its MSI detection accuracy with those of the other three methods on the same samples using the area under receiver operating characteristic curve (AUC). We noticed that even without matched normal samples, MSIsensor-pro’s AUC values were comparable to those of MSIsensor and mantis but much higher than those of mSINGS (Fig. 2a, Supplementary Table 3).

The sequencing data of samples with low coverage and low tumor purities are common challenges for robust MSI detections in clinical applications^17^. To prove the robustness of MSIsensor-pro on various sequencing depths and tumor purities, we evaluated the performances of all four methods on 178 CRC samples (78 MSI and 100 MSS) of both original forms and various sequencing depths, as well as tumor purities. Multiple sequencing depths (5x, 10x, 20x, 40x, 60x and 80x) resulted from simulating and downsampling the original data, while various tumor purities (5%, 10%, 20%, 40%, 60% and 80%) were simulated by mixing the tumor and matched normal samples (Methods). Across samples of diverse depths and tumor purities, the AUCs of MSIsensor-pro, MSIsensor and mantis were all much higher than those of mSINGS, while MSIsensor-pro, requiring only a tumor sample, achieved performance comparable to MSIsenor and mantis, which both required normaltumor-paired samples to call MSI (Fig. 2b, Supplementary Table 4-7). These results confirm the robustness of MSIsensor-pro and indicate MSIsensor-pro can achieve high accuracy on samples with low sequencing depth (e.g. 20x) or low tumor purity (e.g. 40%). In addition, to evaluate the computational performances of all four methods, we called MSI for a TCGA sample TCGA-AD-A5EJ (35Gb tumor and 12Gb normal bam files) using the four methods on a Linux machine running Ubuntu18.04 OS with Intel(R) Core (TM) i5-7500 CPU@3.40 GHz and 32 GB Memory. MSIsensor-pro and MSIsensor required only 4 and 15 minutes, respectively, performances which were significantly faster than mSINGS (94 minutes) and mantis (119 minutes). In addition, MSIsensor-pro consumed much less memory than MSIsensor, mSINGS and mantis (Fig. 2a and Supplementary Fig 11, 12).

While MSIsensor-pro exhibited satisfactory all-around performance in detecting MSI using the 11,666 preselected microsatellites, these sites seemed to have an unequal contribution to the MSI classifications. We therefore evaluated the contribution of each microsatellite based on MND parameter *p* and identified 7,698 sites (Supplementary Table 8) with strong contributions (AUC>0.75), which were defined as discriminative microsatellite (DMS) sites (Supplementary Fig. 13, Supplementary Table 8 and Methods). When DMS sites were used, MSIsensor-pro exhibited slight improvement compared to MSI detection using all 11,666 sites and prevailed over all other methods for the 1,532 TCGA samples. Using DMS sites, MSIsensor-pro’s performance was further enhanced with respect to sequencing data of low depths, especially for depths below 40x (Fig. 2b and Supplementary Table 4-5). For data of different tumor purities, on DMS sites, MSIsensor-pro exhibited performance comparable to other tumor-normal-paired methods for tumor purities of over 40%. However, for lower tumor purities (< 40%), although the performances of all methods decreased, the performance of MSIsensor-pro on DMS sites remained the best of all the examined methods (Fig. 2b and Supplementary Table 6-7).

Since only a portion of all 11,666 sites (DMS sites) was sufficient for high performance MSI calling by MSIsensor-pro, we wondered whether an even smaller subset of DMS sites would be adequate for MSIsensor-pro to achieve similar performance, which shall reduce the time and cost in practical clinical applications. We assessed the MSI calling performance of MSIsensor-pro on microsatellite sets from single and combined tumor samples containing the top 1, 2, 5, 10, 20, 50, 100, 200, 500 and 1,000 DMS sites based on their contributions. We found that even with only 1 top site, MSIsensor-pro had AUC values ranging from 0.92 to 0.96 (Fig. 2c, Supplementary Table 9-10). The performance improved with the increase in top sites and reached a plateau when using the top 20 sites (0.98 AUC). Despite being useful, these top sites would likely suffer from practical issues such as low sequencing coverage. Therefore, by testing the MSIsensor-pro performance on various numbers of randomly selected DMS sites, we sought to identify small panels of DMS sites that are potentially effective enough for robust MSI calling. Indeed, we found that the AUC values of MSI detection steadily increased with growing numbers of randomly selected DMS sites. When as few as 50 random sites were used, the AUC was approximately 0.98 and remained stable. Taken together, MSIsensor-pro could be applied to various target sequencing panels with as few as 50 sites, (Fig. 2d, Supplementary Fig. 14 and Supplementary Table 9-10), indicating MSIsensor-pro’s potential to score MSI with circle tumor DNA or stool DNA. We examined the properties of DMS sites and found that they were closer to splicing sites and located in genes with higher expression than the rest of the sites (Supplementary Fig. 15-17), indicating their potential roles in tumorigenesis.

**Fig. 1.**
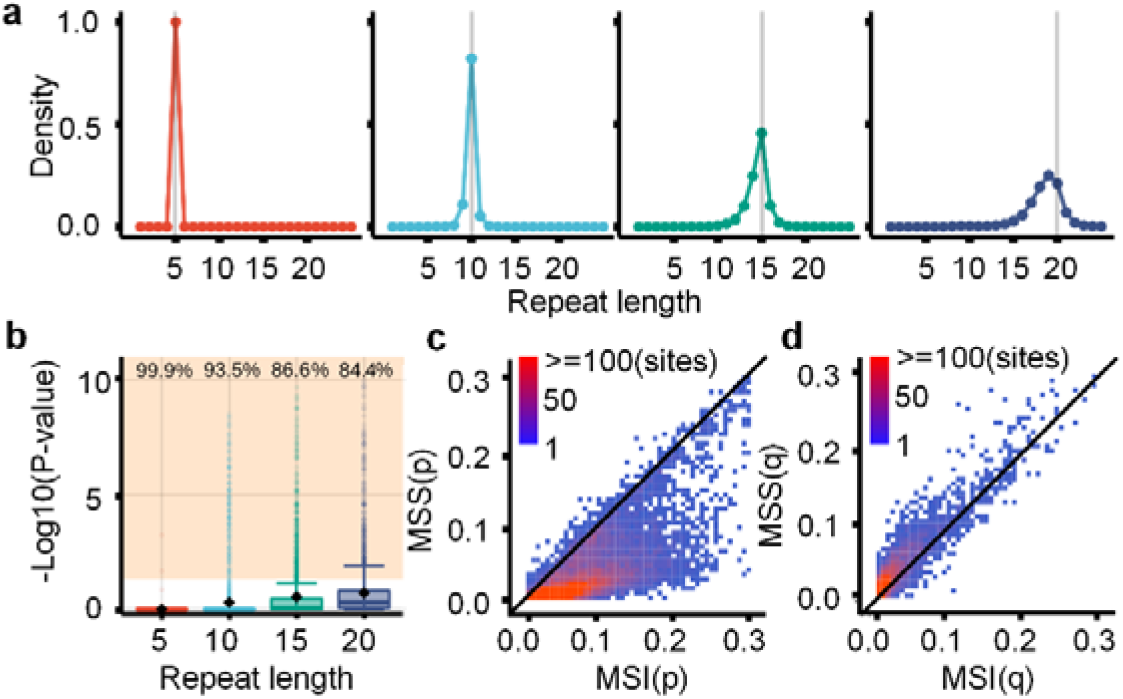
Multinomial distribution model of polymerase slippages. **a**, Allele length distribution of homopolymers in normal samples. The gray vertical lines represent the repeat lengths in the human reference genome (GRCh38). **b**, The fitness of the MND model for polymerase slippages. The points inside the shaded area (*P*-value<0.05) represent sites unfit for the MND model, and the values on the top of boxplot represent the percentage of sites fitted MND model. **c,d,** The dot plots of parameters *p* **(c)** and *q* **(d)** in the MND model between MSI and MSS samples (n=1,532). The color of the points is scaled according to the number of sites. The points near the diagonal lines represent sites undistinguishable between MSI and MSS. MSI samples have significantly larger *p* values than MSS samples **(c)**, while *q* in MSI and MSS was not discriminative **(d)**.

**Fig. 2.**
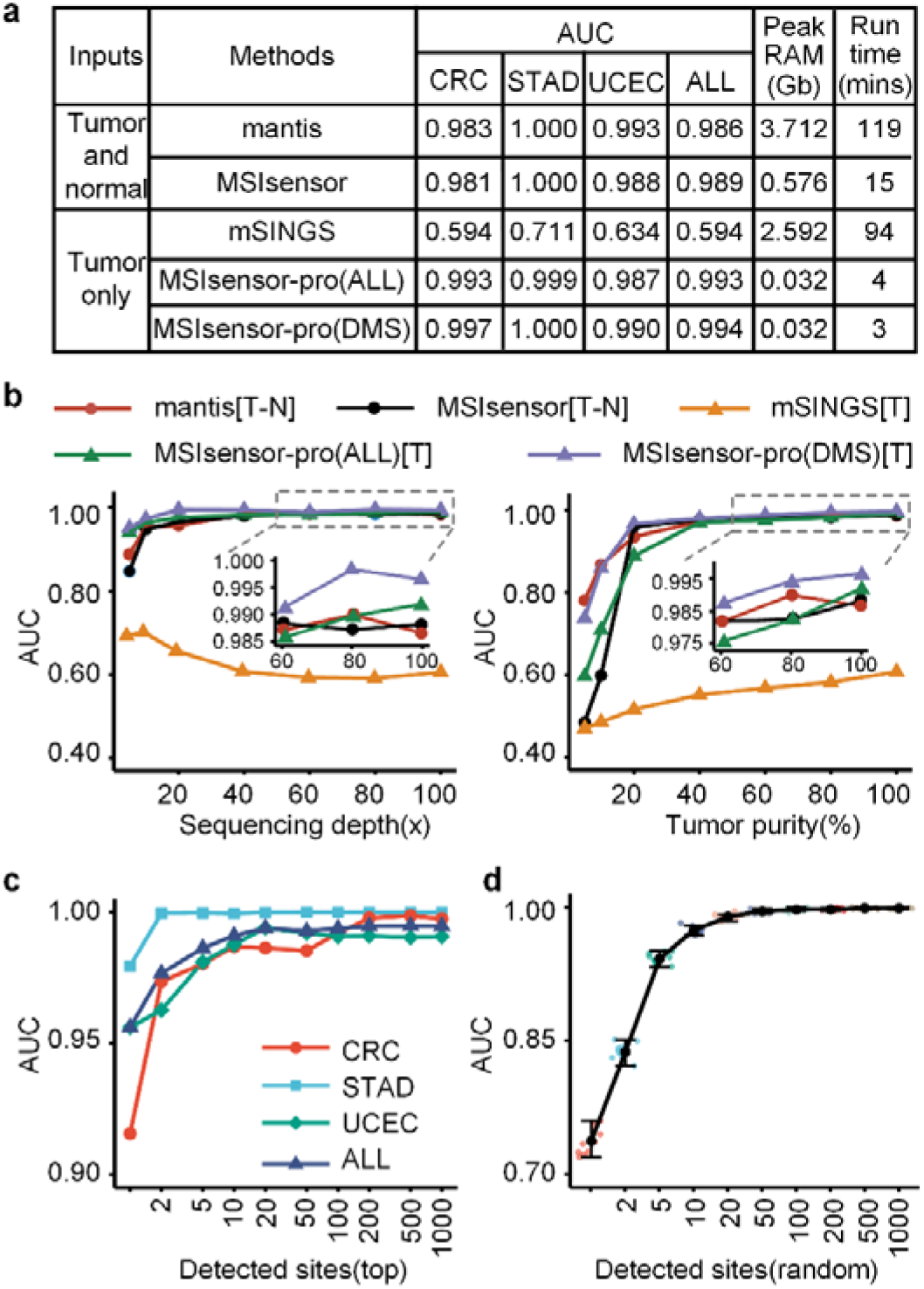
MSI calling accuracy and runtime. **a**, AUCs, peak RAM and runtimes (wallclock time) of four MSI detection methods for 1,532 TCGA samples. **b**, AUCs of four MSI detection methods across various sequencing depths (left) and tumor purities (right) in 78 MSI and 100 randomly selected MSS CRC samples from TCGA. **d,e**, AUCs of MSIsensor-pro with diverse sets of detected sites for 1,532 TCGA samples. The AUCs approach a plateau with the top 20 contributing sites **(d)** and 50 random DMS sites **(e)**. Black points and error bars (random select 10 times) represent the means and standard errors, respectively **(e)**.

## Online Methods

### Data and preprocessing

Whole exome sequencing data and clinical MSI status of a total of 1,532 tumor-normal pairs were downloaded from TCGA^19^ (http://cancergenome.nih.gov/). The sequencing data were aligned against human reference genome (GRCh38), and MSI was determined using the gold standards^20^. The *scan* module (default parameters) in MSIsensor^11^ was used to retrieve the microsatellite regions in the human reference genome. Then, the allele distribution of each microsatellite for each sample was extracted and used in subsequent analyses.

### Multinomial distribution model for polymerase slippage

To detect MSI without matched normal samples, we evaluated the stability of microsatellites using single samples. Based on the characteristics of allele distributions of microsatellites in normal samples (**Supplementary Fig. 1, 2**), we argued that the polymerase slippage during DNA replication is an iterative process and that each step is independently accumulative. Therefore, we use multinomial distribution to model the slippage process in microsatellite sites. We use variable *x* to denote hysteresis synthesis (*x* = 0), presynthesis (*x* = 2) and normal synthesis (*x*=1) of each step of repeat unit synthesis, and the corresponding probabilities are denoted by *p*, *q* and 1–*p–q*, respectively. Then, *x* is subject to a multinoulli distribution, and its probability distribution function is as follows:

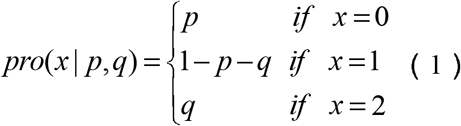

Thus, for a microsatellite site with *n* repeats on the reference genome, we assume that *y* is the repeat length observed from the data. Therefore, we have:

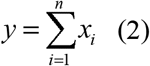

and the probability distribution function of *y* is:

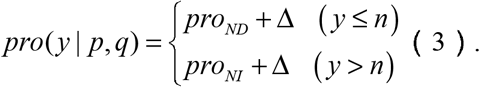

where:

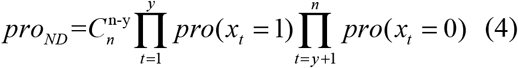

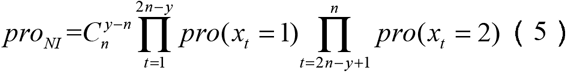

Here, *pro_ND_* and *pro_NI_* denote the probability of acquiring the observed repeat length with minimum steps, while Δ is the probability of using more steps. Since Δ is much smaller and difficult to calculate, we ignore it in practice to save computational resources. For a microsatellite region spanned by *m* reads, we denote the observed repeat length as *y*_1_, *y*_2_,…,*y_t_*…,*y_m_* and its distribution as *Y* = {*y*_1_, *y*_2_,…*y_i_*…,*y_m_*}. Based on *Y*, we use the maximum likelihood estimation to compute *p* and *q* in equation (6).

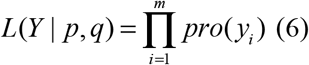

Finally, *p* and *q* can be estimated as follows:

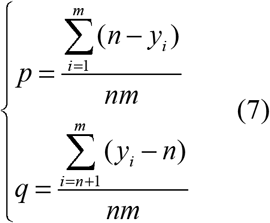

The values of *p* and *q* are positively correlated with the magnitude of polymerase slippages.

### Validation of the MND model

To evaluate how well parameters *p* and *q* in MND mimic polymerases slippage for microsatellites with various repeat lengths, we randomly select 27,200 microsatellites from normal control samples of three cancer types in TCGA and estimate the parameters *p* and *q* of each site. Then, the calculated parameters *p* and *q*, also known as the probabilities of deletion and insertion, are used to simulate allele length distribution. The sites with no significant difference (*P*-value > 0.05, Kolmogorov-Smirnov test) between real and simulated distribution are defined as fitted sites. Then, the percentage of fitted sites to all test sites is used to evaluate the fitness of the MND model. To investigate the polymerase slippages in tumor samples, we estimate *p* and *q* for 1,532 TCGA tumor samples and compare the difference between MSI and MSS samples^5^. We found that *p* is discriminative between the MSI and MSS samples, while *q* is not, indicating that parameter *p* is an effective metric for MSI classification.

### MSI calling of MSIsensor-pro

We use the parameter *p* (probability of deletion) in the MND model to evaluate the stability of microsatellites. To distinguish unstable sites from stable sites, we determine the mean (*μ_i_*) and standard deviation (*σ_i_*) of parameter *p* in the *i*-th microsatellite site in normal samples. Specifically, a microsatellite is classified as unstable when its *p* value is larger than *μ_i_+3σ_i_*. Here, 1,532 normal control samples from three cancer types are used to build the baseline. The MSI score, defined as the percentage of unstable sites within all detected sites in a sample, is used for MSI calling.

### Discriminative microsatellite (DMS) site selection

To find discriminative sites for MSI calling, we compute the contribution of each site to MSI classification. For a given microsatellite site, the parameter *p* is used for MSI classification, and then the AUC is calculated to evaluate the contribution of this site to MSI calling. Finally, the sites with AUC values greater than 0.65 are defined as DMS sites and are used for MSI calling. In this study, 340 TCGA samples are used to discover DMS sites, and the remaining 1,192 samples are used to test the performance of MSIsensor-pro.

### MSIsensor-pro performance evaluation

To assess the performance of MSIsensor-pro, we benchmark MSIsensor-pro against MSIsensor^11^, mantis^15^ and mSINGS^12^ using 1,532 TCGA tumor samples. The MSI score is used for MSI classification, and AUC is used to evaluate the performance of each method (**Supplementary note 1**). The CPU usage, memory and runtime of these methods are tested on a TCGA sample, TCGA-AD-A5EJ, by a Linux machine running Ubuntu18.04 OS with Intel(R) Core (TM) i5-7500 CPU@3.40 GHz and 32 GB memory.

To compare the performances of the four methods on samples with low sequencing depths and low tumor purities, we use 178 CRC tumor-normal paired samples in TCGA to simulate test data. We downsample the raw sequencing data to 5x, 10x, 20x, 40x, 60x and 80x sequencing depths and mix different proportions of tumor and normal sequencing data to generate various tumor purities ranging from 5% to 80%. We call MSI for all simulated data and calculate the AUC for each method. To assess the performance of MSIsensor-pro with fewer sites, we select microsatellite sets containing the top 1, 2, 5, 10, 20, 50, 100, 200, 500, and 1,000 DMS sites for MSI calling. In addition, we randomly select various numbers of microsatellites from DMS sites for MSI calling to examine how many sites are sufficient for MSI calling by MSIsensor-pro.

## Code availability

MSIsensor-pro source code is freely available at https://github.com/xjtuomics/msisensor-pro with help documentation and demo data for testing.

## Data availability

Primary sequencing data, gold standard MSI status and RNA expression data are downloaded from TCGA Research Network (http://cancergenome.nih.gov/). All results generated by this study are available within the article and in the Supplementary Data, or are available from the authors upon request.

## Acknowledgments

We thank T. Wang, Y. Kang, X. Li and S. Gao for helpful discussions regarding data analysis and J. Hai for administrative and technical support. This study was supported by the National Key R&D Program of China (grant Nos. 2018YFC0910400 and 2017YFC0907500), the National Science Foundation of China (grant Nos. 31671372, 61702406, 31701739 and 31970317), the National Science and Technology Major Project of China (grant No. 2018ZX10302205), the “World-Class Universities and the Characteristic Development Guidance Funds for the Central Universities” and the General Financial Grant from the China Postdoctoral Science Foundation (2017M623178 and 2017M623188).

## Author contributions

K.Y conceived of, designed and supervised the study; P.J and B.L developed the multinomial distribution model for polymerase slippage estimation; P.J and H.L implemented the source code of MSIsensor-pro; P.J evaluated the performances of MSIsensor-pro and the other four MSI detection methods. P.J, X.Y, L.G and K.Y wrote the manuscript, and all authors contributed to the critical revision of the manuscript and approved the final version.

