## Supplementary material for "MSIsensor-pro: fast, accurate and matched-normal-sample-free detection of microsatellite instability": Suplemental text

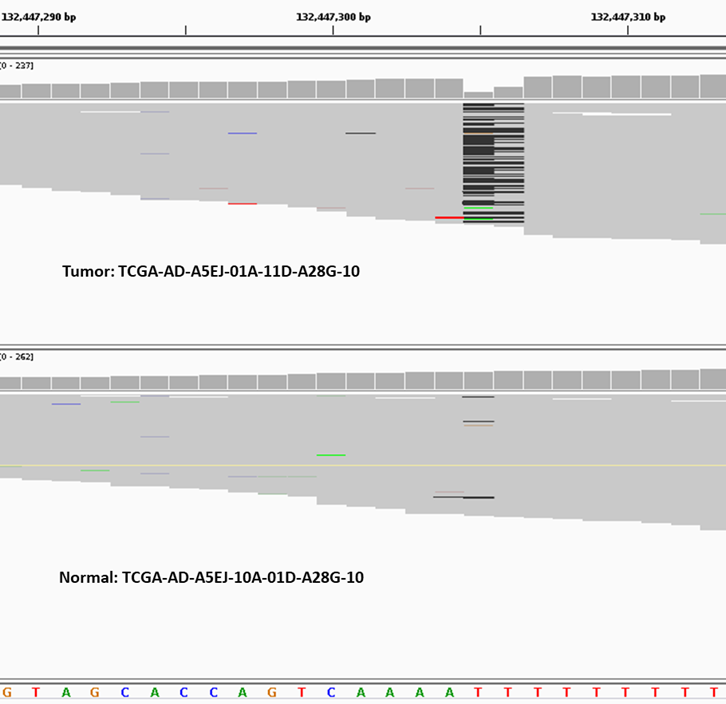

Supplementary Figure 1.

The difference characteristics of an MSI case between a tumor sample and matched normal sample.

This IGV screenshot of the sample with MSI-H status (TCGA-AD-A5EJ) shows that chr3_132447304 microsatellites contain more deletions in the tumor sample than in the matched normal sample.

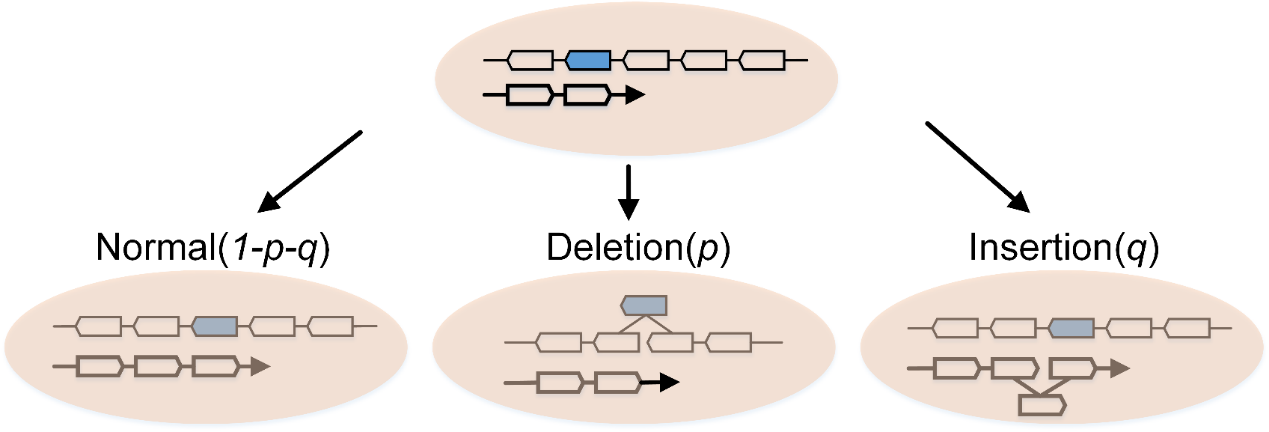

Supplementary Figure 2.

Basic model for polymerase slippage.

When the DNA strand in the top panel continues to synthesize, there are three possibilities for the next step: normal, deletion and insertion. This process can be described as a multinoulli distribution.

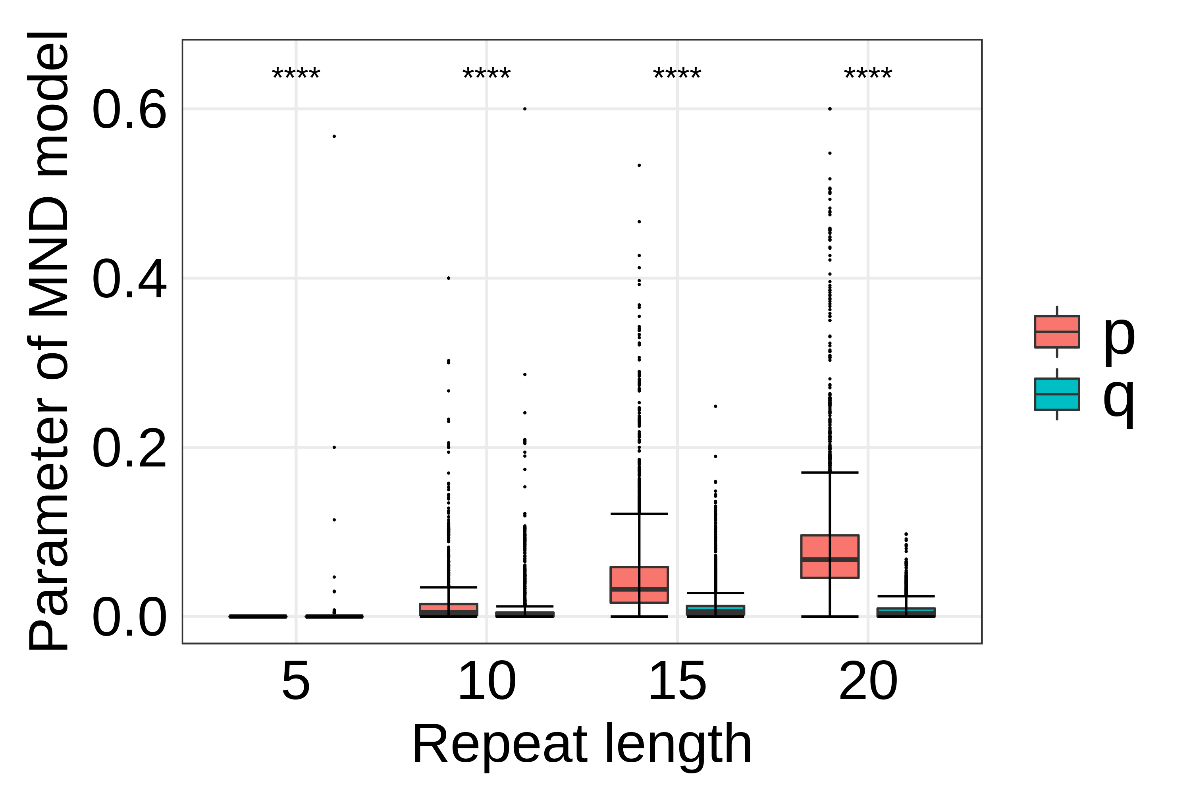

Supplementary Figure 3.

Boxplot of parameters *p* and *q* in the MND model.

The boxplot shows that the polymerase slippages are accumulative with the increase in the microsatellite repeat length. The values of parameter *p* in test microsatellites are significantly larger than those of parameter *q*. Rank-sum test are implemented between MSI and MSS samples. ns: p>0.05; *: p<0.05; **: p<0.01; ***: p<0.001; ****: p<0.0001.

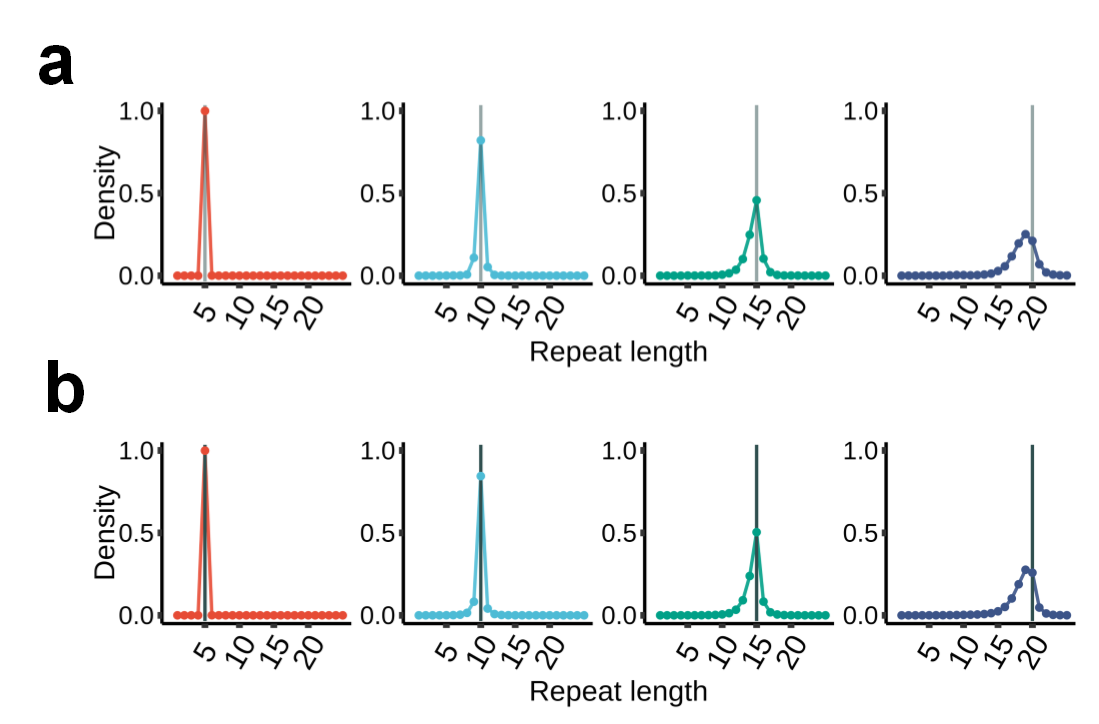

Supplementary Figure 4.

Allele length distributions of homopolymers.

**a.** The average allele length distributions of homopolymers with 5, 10, 15 and 20 repeats. **b.** The simulated allele length distributions from the calculated p (probability deletion) and q (probability insertion).

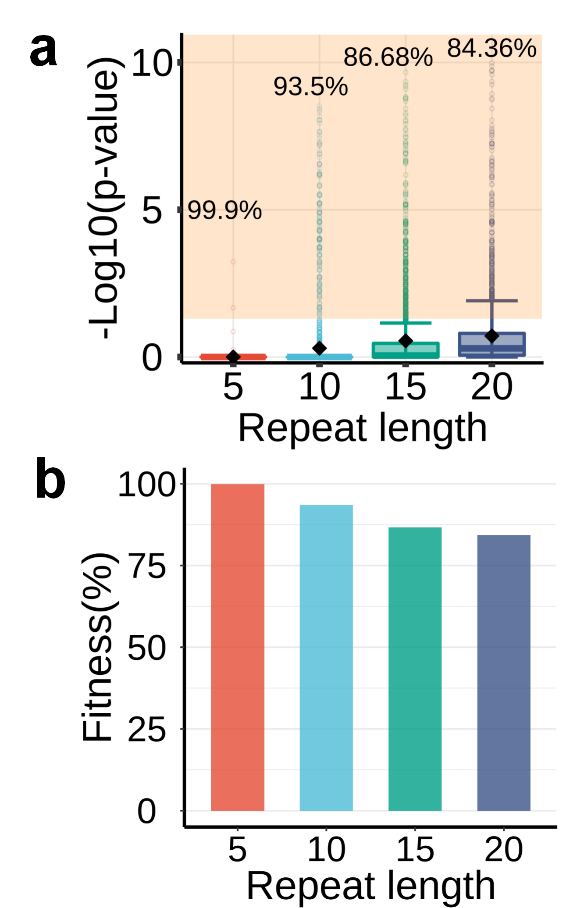

Supplementary Figure 5.

Performance of MND for polymerase slippage estimation in TCGA normal samples.

**a.** p-values of Kolmogorov-Smirnov testing between the observed allele distribution and simulated results; the background represents p-values less than 0.05. **b.** Fitness of MND for polymerase slippages, represented by the percentage of sites with p-values less than 0.05 in panel **(a).**

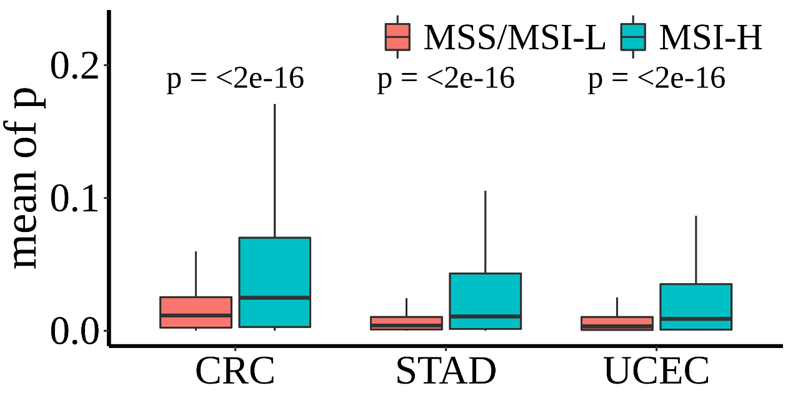

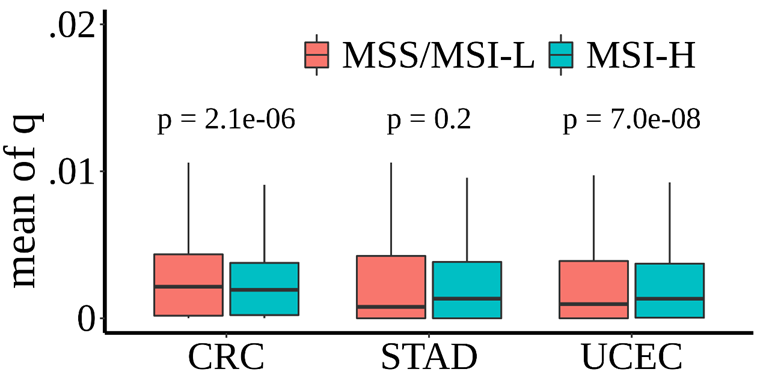

Supplementary Figure 6.

Parameters p and q of the MND model in different MSI status samples.

The boxplot shows the mean of p (top panel) and q (bottom panel) for each site in MSI-P and MSI-N samples in the three tumors. Rank-sum tests are implemented between MSI and MSS samples.

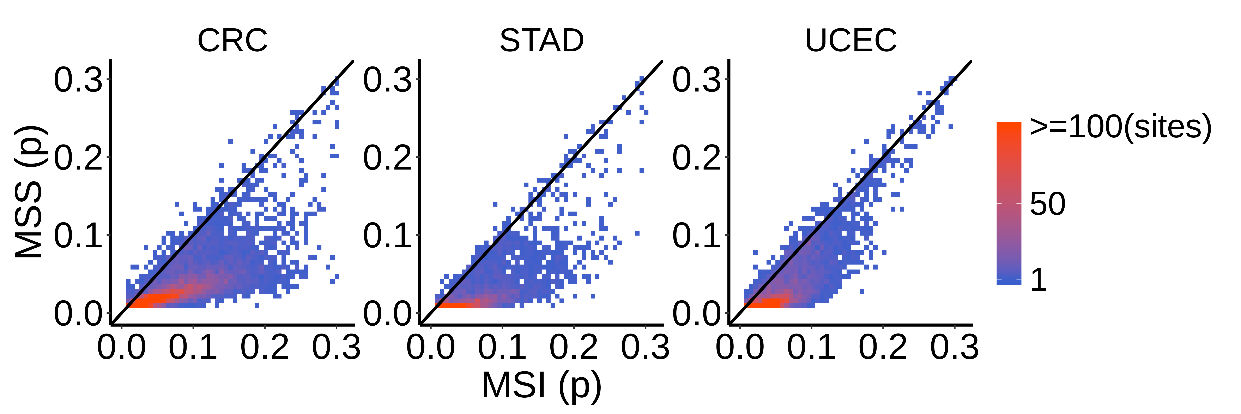

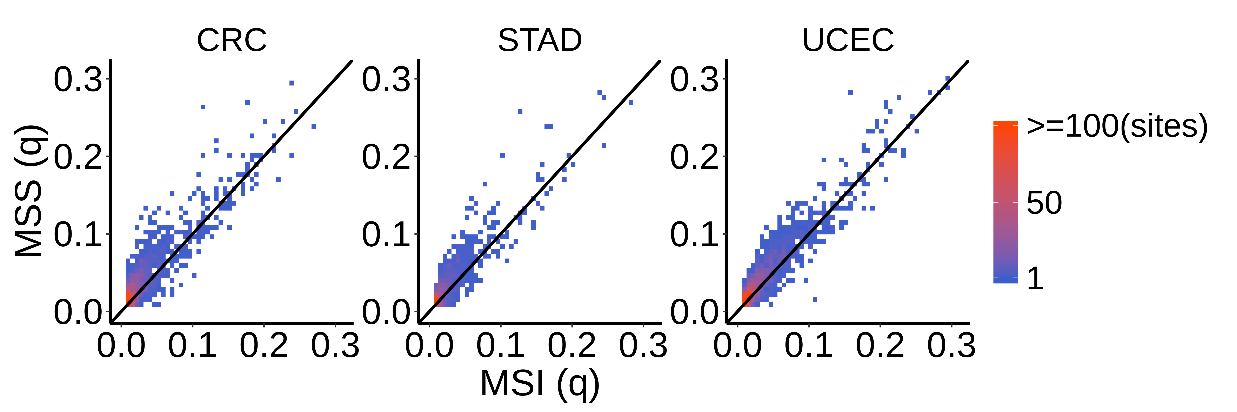

Supplementary Figure 7.

Difference of parameters p and q of the MND model between MSI and MSS samples.

The horizontal and vertical coordinates represent the means of PSM in MSI-P and MSI-N, respectively. Most of the p (top) values in MSI are larger than those in MSS, while there is no such result for q (bottom). The colors of sites represent the density of sites.

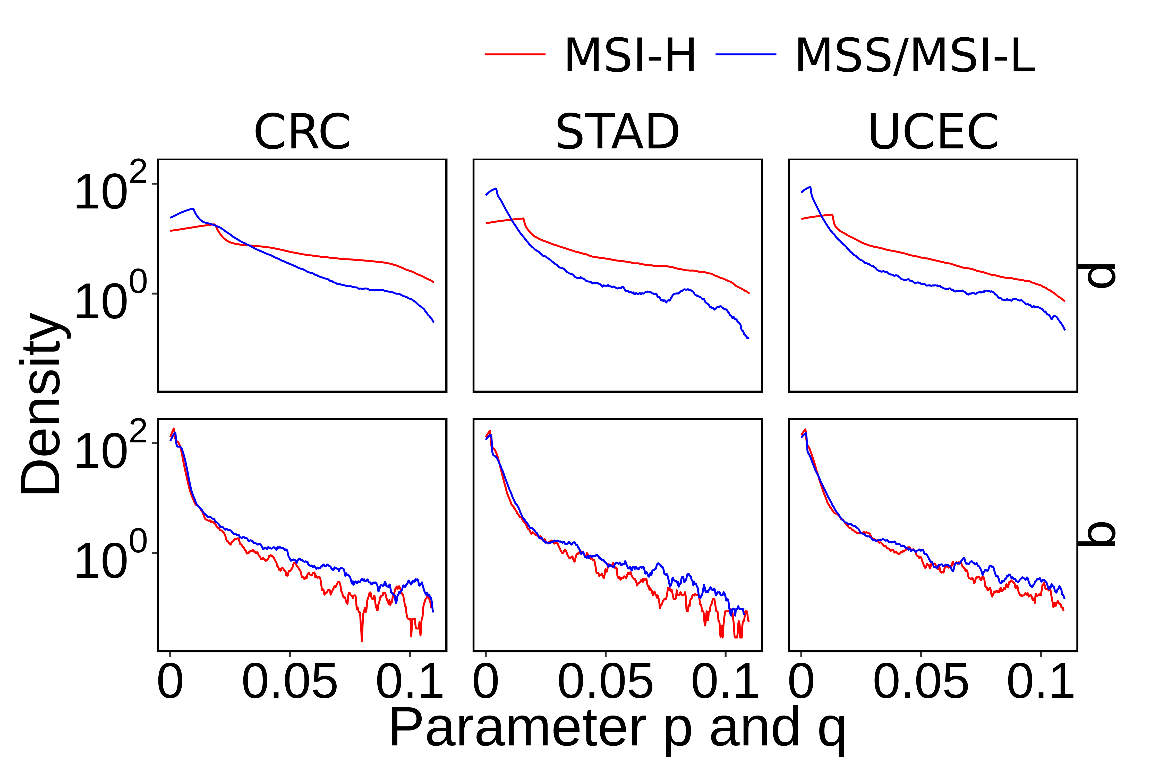

Supplementary Figure 8.

Density plot of parameters p and q in TCGA samples.

The values of horizontal coordinates for each site are calculated by the mean of p and q in different MSI status and different tumor types. The results show that MSI samples have more sites with larger p values.

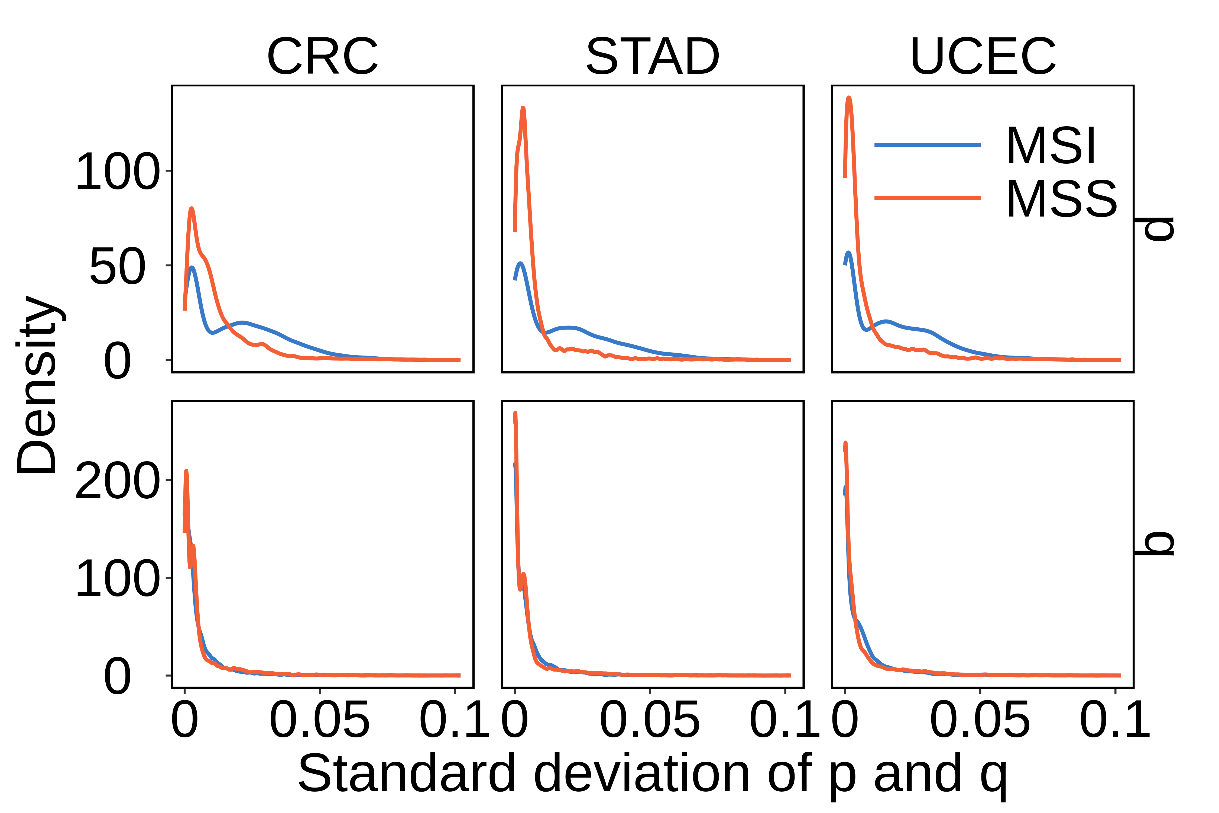

Supplementary Figure 9.

Density plots for the standard deviations of parameters p and q in TCGA samples

The density plots of standard deviations of p (top panel) and q (bottom panel) in three types of tumors. The values of horizontal coordinates for each site are calculated by standard deviation of p and q in different MSI status and different tumor types.

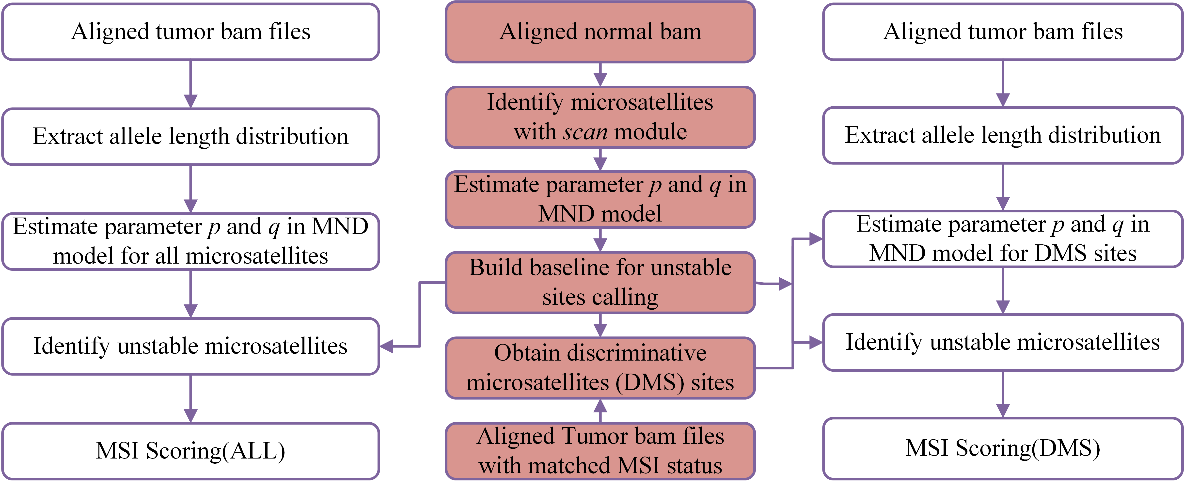

Supplementary Figure 10.

Workflow of MSIsensor-pro.

The processes with red background are basic processes for baseline building and discriminative microsatellites (DMS) site selection. The left processes show the MSIsensor-pro score of MSI for all sites, whereas the right shows DMS sites.

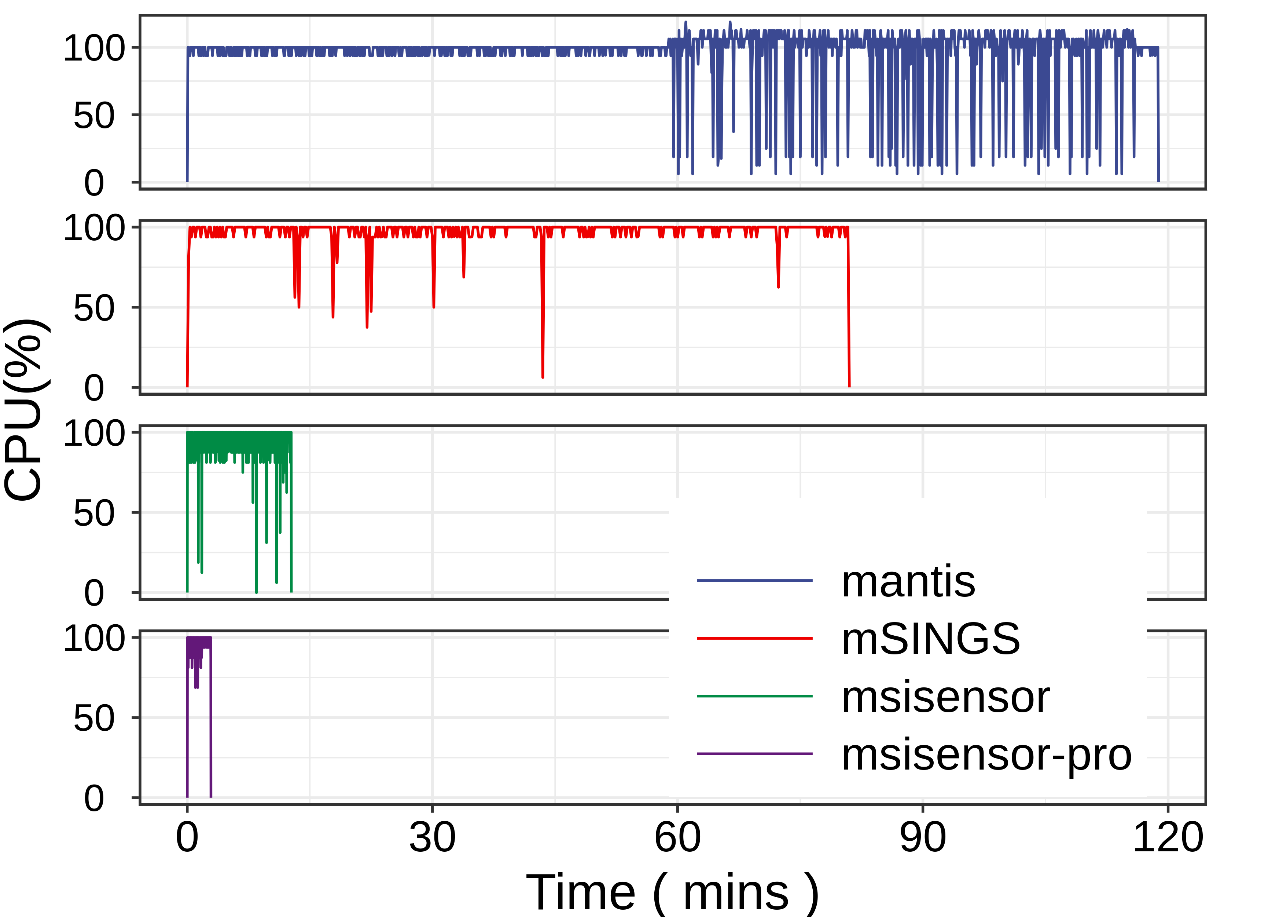

Supplementary Figure 11.

CPU usage along with runtime for MSI calling methods.

CPU usage and runtime are tested by running TCGA-AD-A5EJ using MSIsensor-pro and three other methods on Ubuntu18.04 OS with an Intel(R) Core (TM) i5-7500 CPU@3.40 GHz and 32 GB memory.

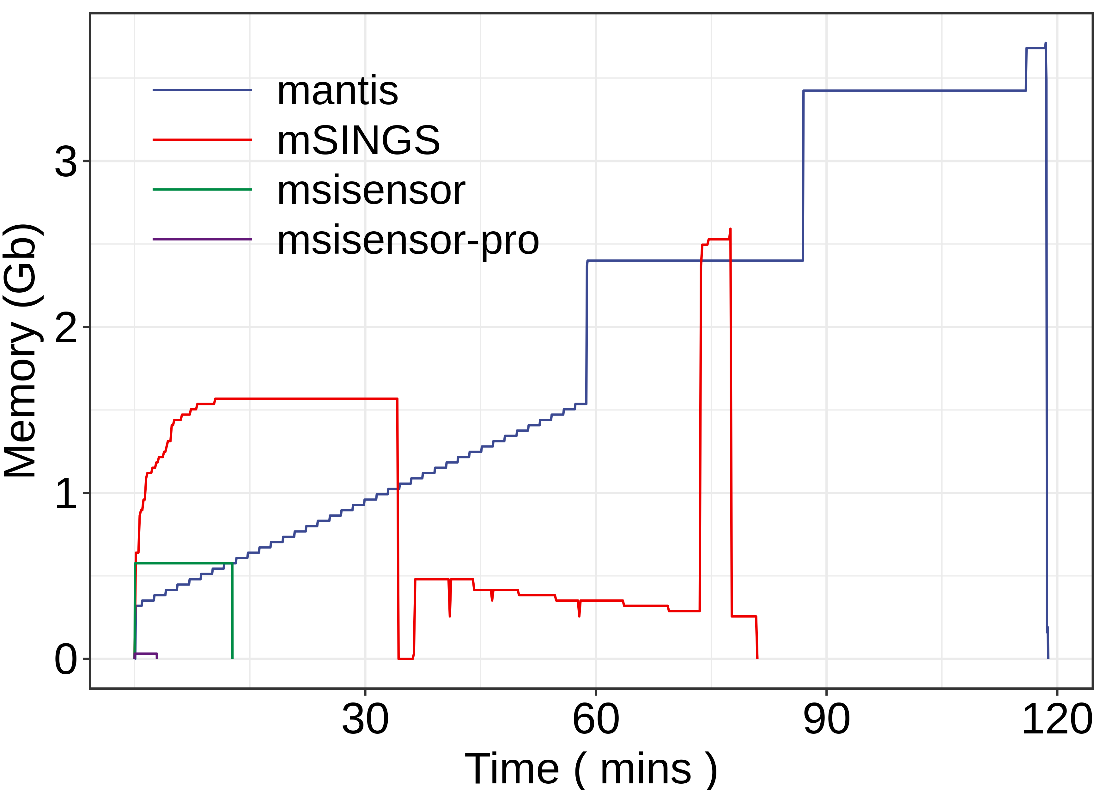

Supplementary Figure 12.

Memory usage along with runtime for MSI calling methods.

Memory usage and runtime are tested by running TCGA-AD-A5EJ using MSIsensor-pro and 3 other methods on Ubuntu18.04 OS with an Intel(R) Core (TM) i5-7500 CPU@3.40 GHz and 32 GB memory.

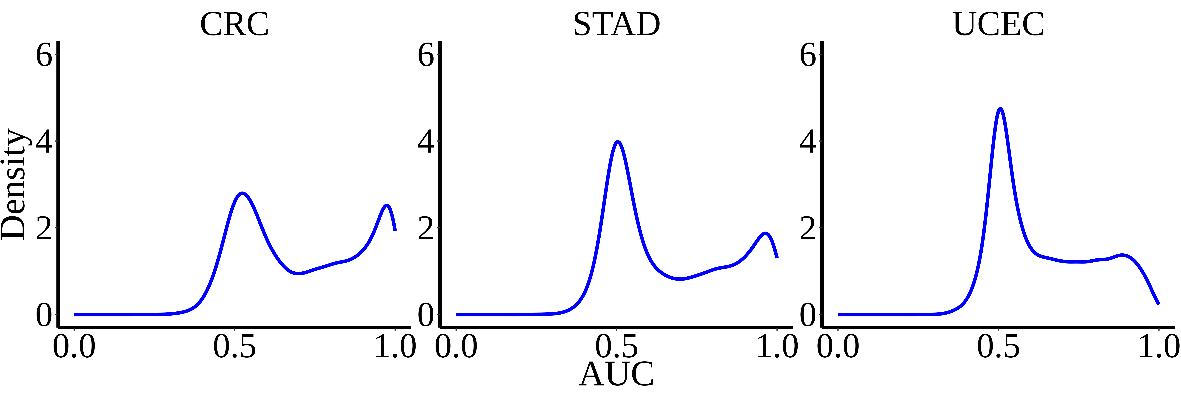

Supplementary Figure 13.

Density plots of site contributions for MSI calling.

The contribution of each site is calculated by its AUC for MSI calling.

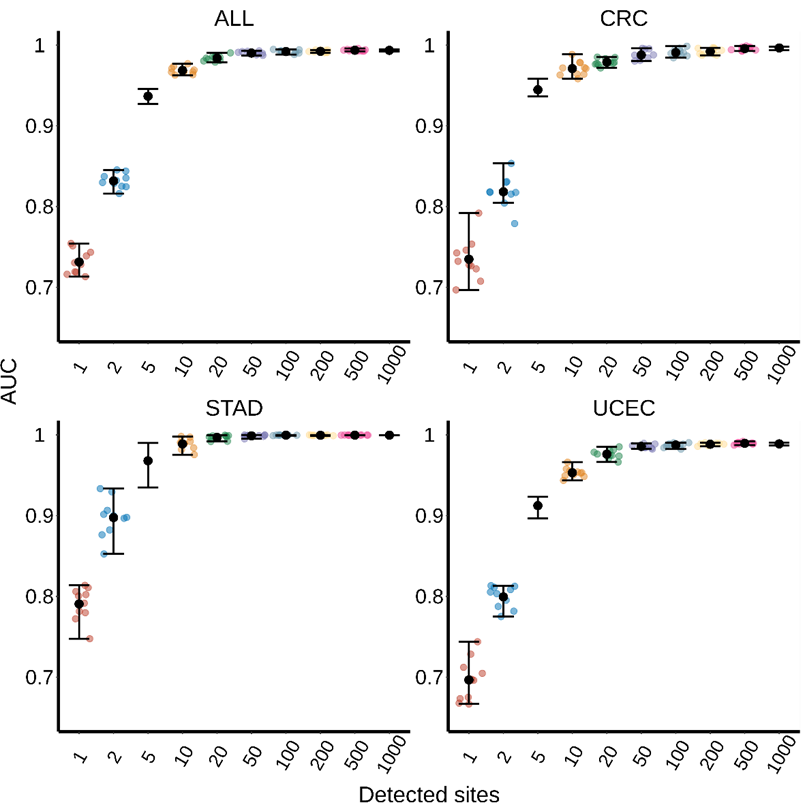

Supplementary Figure 14.

Performance of MSIsensor-pro for different site sets.

Here, we randomly select 1, 2, 5, 10, 20, 50, 100, 200, 500, and 1,000 DMS sites for MSI calling using MSIsensor-pro. These random tests are implemented 10 times. Each color point represents the AUC of one random test, the black point is the mean of 10 AUC values, and the top line and bottom lines of each bar are the maximum and minimum of 10 AUCs.

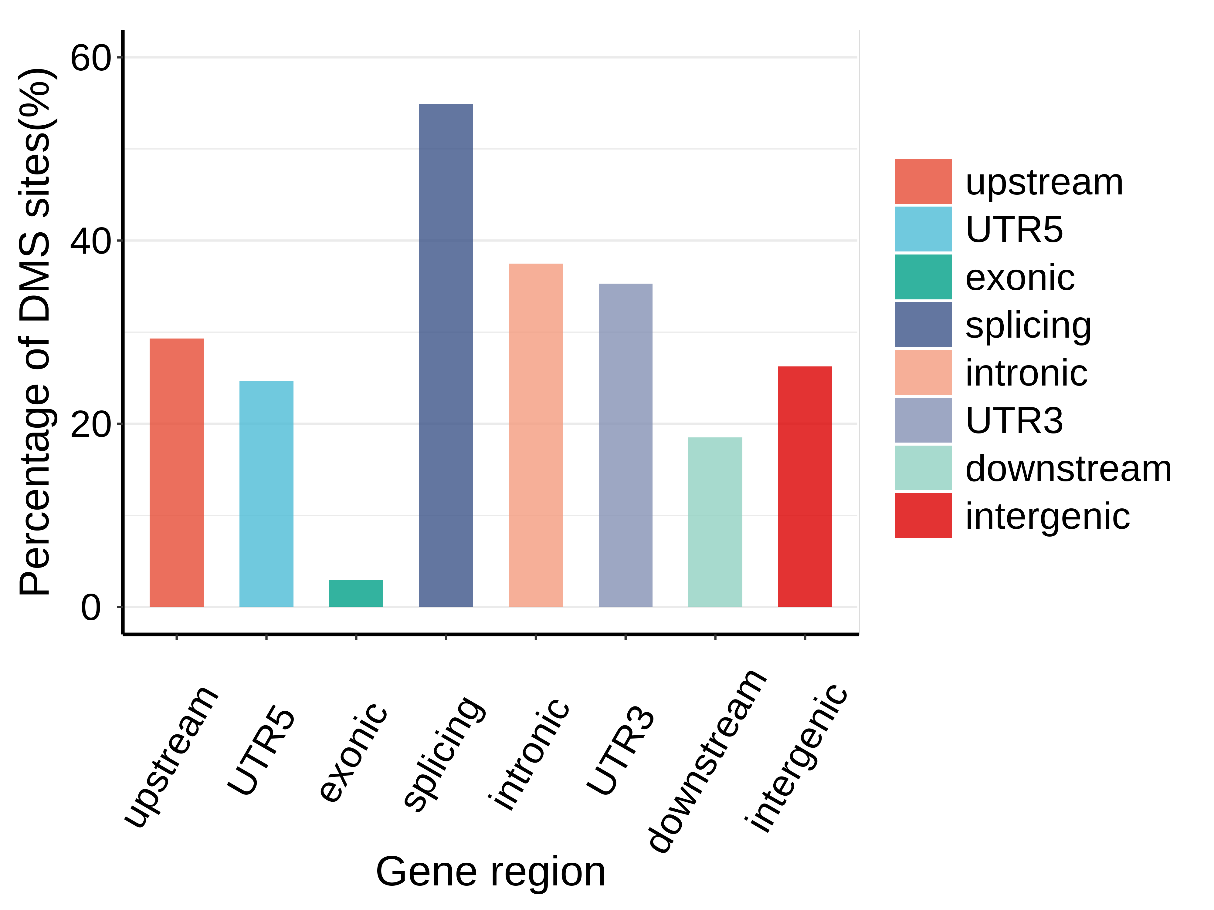

Supplementary Figure 15.

The percentage of DMS sites in each gene region.

The bar plot shows that there are more DMS sites in covered splicing regions and fewer in exonic regions.

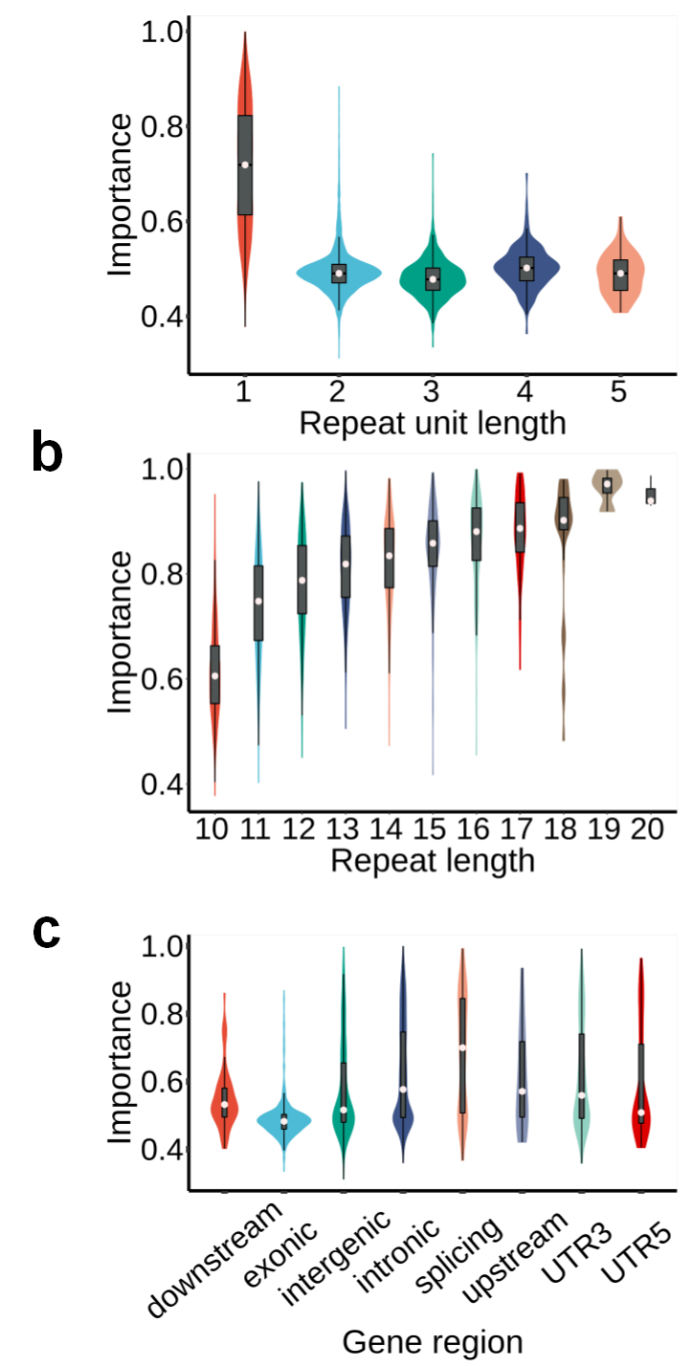

Supplementary Figure 16.

The average importance of sites by different repeat unit length, repeat length and gene region.

a. Homopolymers have greater contributions to MSI classification than sites with more than 2 repeat units. **b.** For homopolymers, the contributions increase with increasing repeats. **c.** The microsatellites covering splicing sites have greater contribution to MSI classification than other regions.

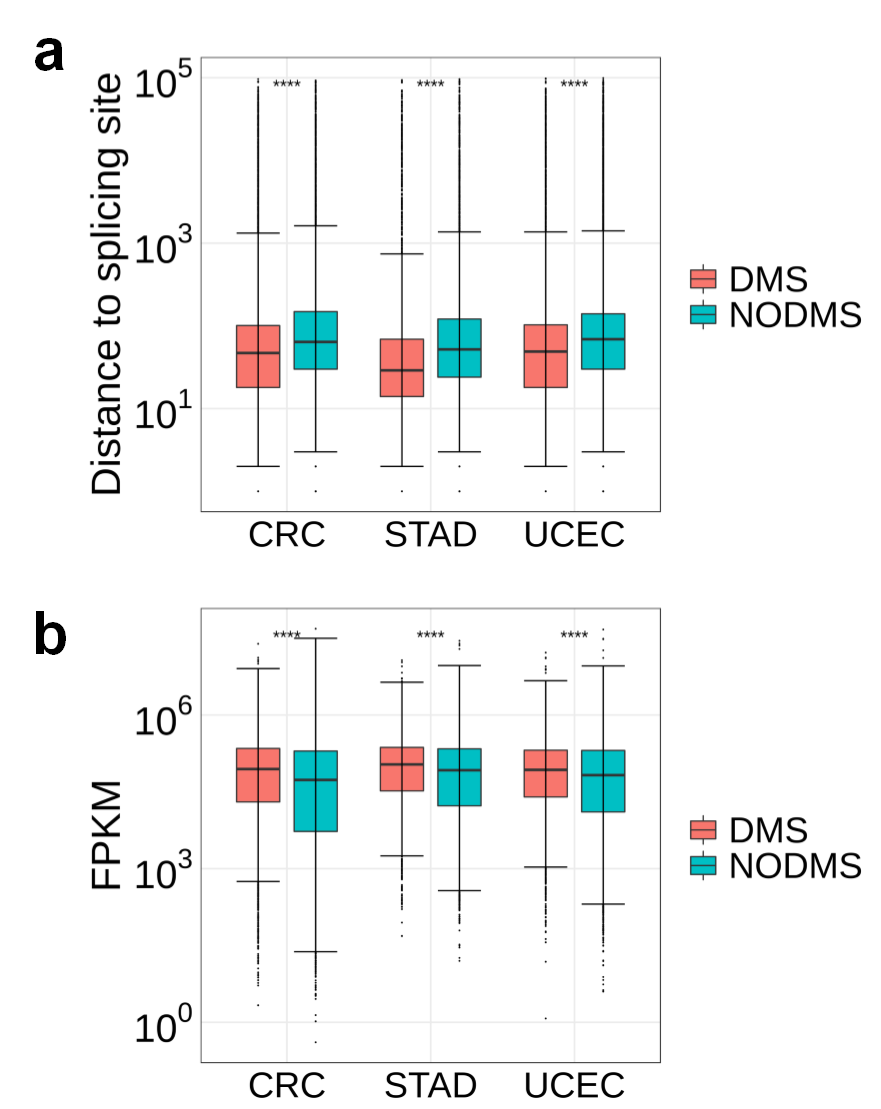

Supplementary Figure 17.

The characteristics of DMS sites.

**a.** The DMS sites are closer to the splicing sites than the remaining sites. **b.** The genes containing DMS sites exhibit higher expression than the genes covered remaining sites. Rank-sum tests are implemented between DMS sites and the remaining sites. ns: p>0.05; *: p<0.05; **: p<0.01; ***: p<0.001; ****: p<0.0001.

**Supplementary Table 1: Overview of MSI status in TCGA samples**

| Cancer types | MSI-H | MSS/MSI-L | Total |
| --- | --- | --- | --- |
| CRC | 28/78 | 109/510 | 137/588 |
| STAD | 28/80 | 69/332 | 97/412 |
| UCEC | 59/168 | 47/364 | 106/532 |
| Total | 115/326 | 225/1206 | 340/1532 |

*N/M: M is the sample number, N is the number of the sample for the importance determination of microsatellites.

**Supplementary Table 5: MSI calling AUCs of low sequencing depth data**

| Sequencing depths(x) | mSINGS | mantis | MSIsensor | MSIsensor-pro(ALL) | MSIsensor-pro(DMS) |
| --- | --- | --- | --- | --- | --- |
| 5 | 0.6932 | 0.8902 | 0.8491 | 0.9425 | 0.9535 |
| 10 | 0.7022 | 0.9553 | 0.9500 | 0.9657 | 0.9753 |
| 20 | 0.6575 | 0.9592 | 0.9678 | 0.9786 | 0.9977 |
| 40 | 0.6085 | 0.9868 | 0.9824 | 0.9849 | 0.9971 |
| 60 | 0.5938 | 0.9871 | 0.9883 | 0.9856 | 0.9912 |
| 80 | 0.5923 | 0.9899 | 0.9872 | 0.9897 | 0.9983 |
| 100 | 0.6062 | 0.9866 | 0.9882 | 0.9919 | 0.9965 |

**Supplementary Table 7: MSI calling AUCs of low tumor purity data**

| Tumor purities (%) | mSINGS | mantis | MSIsensor | MSIsensor-pro(ALL) | MSIsensor-pro(DMS) |
| --- | --- | --- | --- | --- | --- |
| 5 | 0.4718 | 0.7807 | 0.4853 | 0.5971 | 0.7378 |
| 10 | 0.4873 | 0.8679 | 0.5991 | 0.7088 | 0.8596 |
| 20 | 0.5172 | 0.9347 | 0.9597 | 0.8885 | 0.9664 |
| 40 | 0.5528 | 0.9724 | 0.9767 | 0.9701 | 0.9797 |
| 60 | 0.5690 | 0.9816 | 0.9819 | 0.9755 | 0.9872 |
| 80 | 0.5831 | 0.9899 | 0.9823 | 0.9822 | 0.9942 |
| 100 | 0.6079 | 0.9866 | 0.9882 | 0.9919 | 0.9965 |

**Supplementary Table 2: Detailed information of 1,532 TCGA samples**

**Supplementary Table 3: MSI calling AUCs of MSIsensor-pro and three other MSI detection methods**

**Supplementary Table 4: MSI calling results of low sequencing depth data**

**Supplementary Table 6: MSI calling results of low tumor purity data**

**Supplementary Table 8: Microsatellite site information**

**Supplementary Table 9: MSI calling results of selected DMS sites**

**Supplementary Table 10: MSI calling AUCs of selected DMS sites**

**Supplementary note 1:**

To assess the performance of MSIsensor-pro, we applied MSIsensor-pro, MSIsensor, mantis, and mSINGS to TCGA samples. The following are the versions of each software and the parameter details.

**MSIsensor-pro:**

- Address: <https://github.com/xjtu-omics/msisensor-pro>
- Version: v0.1.0
- Microsatellites information: scan module
- Minimum read coverage(-c): 5 for sequencing coverage less than 10x
- Other parameters: default

**MSIsensor:**

- Address: <https://github.com/ding-lab/msisensor>
- Version: v0.1.0
- Microsatellites information: scan module
- Minimum read coverage(-c): 5 for sequencing coverage less than 10x
- Other parameters: default

**mantis:**

- Address: <https://github.com/OSU-SRLab/MANTIS>
- Version: v1.0.4
- Microsatellites information: RepeatFinder
- Minimum read coverage (-mlc): 5 for sequencing coverage less than 10x
- Other parameters: default

**mSINGS:**

- Address: <https://bitbucket.org/uwlabmed/msings/src/master/>
- Version: 0191.0302893
- Microsatellites information: MISA
- Other parameters: default
- Notes: The baseline was built according to the instructions on the mSINGS website, and the same samples were used in building the baseline.

**Supplementary note 2:** Main commands of MSIsensor-pro

1. Baseline:

| baseline | | |
| --- | --- | --- |
| Function | This module builds the baseline for MSI detection with the pro module using only tumor sequencing data. To achieve this goal, sequencing data from normal samples is required (-i). | |
| Example | msisensor-pro baseline -d /path/to/reference.list -i \  /path/to/configure.txt -o /path/to/baseline/directory | |
| Parameter | Type | Explanation |
| -d | <string> | homopolymer and microsatellite file |
| -i | <string> | configure files for building baseline (text file) |
| -o | <string> | output directory |
| -c | <int> | coverage threshold for msi analysis, WXS: 20; WGS: 15, default=20 |
| -i | <double> | a site with a detected ratio in all samples less than this parameter will be removed in following analysis, default=0.5 |
| -p | <int> | minimal homopolymer size for pro analysis, default=10 |
| -m | <int> | maximal homopolymer size for pro analysis, default=50 |
| -u | <int> | span size around window for extracting reads, default=500 |
| -s | <int> | minimal microsatellite size for distribution analysis, default=5 |
| -w | <int> | maximal microsatellite size for distribution analysis, default=40 |
| -b | <int> | thread number for parallel computing, default=1 |
| -x | <int> | output homopolymer only, 0: no; 1: yes, default=0 |
| -y | <int> | output microsatellite only, 0: no; 1: yes, default=0 |
| -0 | <int> | output site with no read coverage, 1: no; 0: yes, default=0 |
| -h | -- | help |

2. pro:

| pro | | |
| --- | --- | --- |
| Function | This module evaluates MSI using tumor only samples. Required inputs are (-d) microsatellites file and bam files (-t). | |
| Example | 1. msisensor-pro pro -d /path/to/reference.list -i 0.1 -t \   /path/to/case1_tumor_sorted.bam -o /path/to/case1_output   2. msisensor-pro pro -d /path/to/reference.list_baseline –t\   /path/to/case1_tumor_sorted.bam -o /path/to/case1_output | |
| -d | <string> | homopolymer and microsatellites file |
| -t | <string> | tumor bam file |
| -o | <string> | output prefix |
| -e | <string> | bed file, optional |
| -i | <double> | minimal threshold for unstable site detection (for tumor only data), default=0.1 |
| -c | <int> | coverage threshold for msi analysis, WXS: 20; WGS: 15, default=20 |
| -r | <string> | choose one region, format: 1:10000000-20000000 |
| -p | <int> | minimal homopolymer size for distribution analysis, default=10 |
| -m | <int> | maximal homopolymer size for distribution analysis, default=50 |
| -s | <int> | minimal microsatellite size for distribution analysis, default=5 |
| -w | <int> | maximal microsatellite size for distribution analysis, default=40 |
| -u | <int> | span size around window for extracting reads, default=500 |
| -b | <int> | thread number for parallel computing, default=1 |
| -x | <int> | output homopolymer only, 0: no; 1: yes, default=0 |
| -y | <int> | output microsatellite only, 0: no; 1: yes, default=0 |
| -0 | <int> | output site with no read coverage, 1: no; 0: yes, default=0 |
| -h |  | Help |
